# Rotavirus Replication Factories Are Complex Ribonucleoprotein Condensates

**DOI:** 10.1101/2020.12.18.423429

**Authors:** Florian Geiger, Guido Papa, William E. Arter, Julia Acker, Kadi L. Saar, Nadia Erkamp, Runzhang Qi, Jack Bravo, Sebastian Strauss, Georg Krainer, Oscar R. Burrone, Ralf Jungmann, Tuomas P.J. Knowles, Hanna Engelke, Alexander Borodavka

## Abstract

RNA viruses induce formation of subcellular organelles that provide microenvironments conducive to their replication. Here we show that replication factories of rotaviruses represent protein-RNA condensates that are formed via liquid-liquid phase separation. We demonstrate that rotavirus proteins NSP5 and NSP2 undergo phase separation *in vitro* and form RNA-rich condensates *in vivo* that can be reversibly dissolved by aliphatic diols. During infection, these RNA-protein condensates became less dynamic and impervious to aliphatic diols, indicating a transition from a liquid to solid state. Some aspects of assembly of rotavirus replication factories mirror the formation of cytoplasmic ribonucleoprotein granules, while the selective enrichment of viral transcripts appears to be a unique feature of these condensates. Such complex RNA-protein condensates that underlie replication of RNA viruses represent an attractive target for developing novel therapeutic approaches.

## Introduction

To replicate successfully, RNA viruses compartmentalise their replicative enzymes within specialized organelles termed viral factories. These structures are viewed as virus assembly lines that support viral replication by sequestering and concentrating cognate nucleic acids and proteins. While most viral RNA replication requires membrane-enclosed replication compartments, experimental evidence from recent studies^1–4^ suggests that liquid–liquid phase separation (LLPS) may provide a simple solution to assembling RNA-rich replication factories via a process that is solely dependent upon physical forces^5–10^.

Here we show that replication factories of rotaviruses (RVs), a large class of human and animal double-stranded RNA (dsRNA) viruses, are formed via phase separation of the viral non-structural proteins NSP5 and NSP2. Both proteins are indispensable for rotavirus replication, constituting the bulk of the replication factories, or viroplasms^11–16^. In RV-infected cells, large amounts of NSP5 and NSP2 rapidly accumulate in the cytoplasm, forming viroplasms as early as 2 hours post infection^12,17,18^. We demonstrate that upon mixing at low micromolar concentrations *in vitro*, or when co-expressed in cells, multivalent Ser/Glu-rich protein NSP5 and the viral RNA chaperone NSP2 spontaneously phase separate forming liquid condensates. Analysis of replication of the NSP5-defficient recombinant rotavirus in a cell line stably expressing NSP5 confirms that the condensate formation requires NSP5. Both rotavirus replication factories and NSP5/NSP2 condensates were rapidly and reversibly dissolved in the presence of small aliphatic alcohols, including 1,6-hexanediol, as well as lower molecular weight propylene diols, corroborating their liquid-like properties. We have validated our findings by employing a combinatorial droplet microfluidic platform, termed PhaseScan^19^, to characterise the phase behaviour of the NSP5/NSP2 condensates and mapped out the phase boundary, at which they transition from a mixed one-phase, to a two-phase demixed state.

Finally, using single-molecule RNA fluorescence in situ hybridization (smFISH) and super-resolution DNA-PAINT imaging, we have shown that reversible dissolution of replication factory condensates releases rotavirus transcripts, followed by their reassociation upon removal of these compounds. At a later infection (>12 h) stage, the apparent shapes of rotavirus replication factories deviated from a perfect sphere, and did not dissolve in the presence of aliphatic diols. These observations were consistent with the decreased exchange of NSP5-EGFP between the cytoplasm and viroplasm during late infection, suggesting a liquid-to-solid transition that occurs during infection.

The emerging properties of these viral protein–RNA condensates in a large family of dsRNA viruses are remarkably similar to recent results emerging from studies of non-membrane-bound cytosolic ribonucleoprotein (RNP) organelles, including Processing (P) bodies and stress granules. Their capacity to rapidly and reversibly respond to external stimuli amounts to a shift in our understanding of the replication of multi-segmented viral RNA genomes, providing the basis for viewing these RNA–protein condensates as an attractive target for developing novel antiviral therapeutics.

## Results

### Liquid-like Properties of Rotavirus Replication Factories

The dynamic nature of the RNA-rich viral cytoplasmic inclusions previously termed ‘viroplasms’, and their tendency to coalesce^17,18^ during rotavirus (RV) infection are reminiscent of other cytoplasmic liquid-like ribonucleoprotein cytosolic granules^20^. Such observations have prompted us to further investigate the liquid-like properties of viroplasms.

Previous reports demonstrated that the two viral proteins NSP5 and NSP2 constitute the bulk of viroplasms^13,17,21–23^. We used MA104 cell lines that fully support RV replication, whilst expressing low levels of the C-terminally EGFP- and mCherry-tagged NSP5 and NSP2, respectively^16,17^. Upon RV infection, both cytosolic NSP2-mCherry^16^ and NSP5-EGFP^18^ relocalise into newly formed replication factories, thus making them suitable markers for live-cell imaging of these virus-induced organelles.

At 4 hours post infection (HPI), more than 90% of virus-infected NSP5-EGFP or NSP2-mCherry cell lines contained NSP5-EGFP or NSP2-mCherry-containing cytoplasmic granules, respectively. We were able to observe fusion events between these granules, irrespective of the fusion fluorescent reporter protein used (**Fig. 1a**), suggesting that these inclusions may have liquid-like properties.

**Figure 1.**
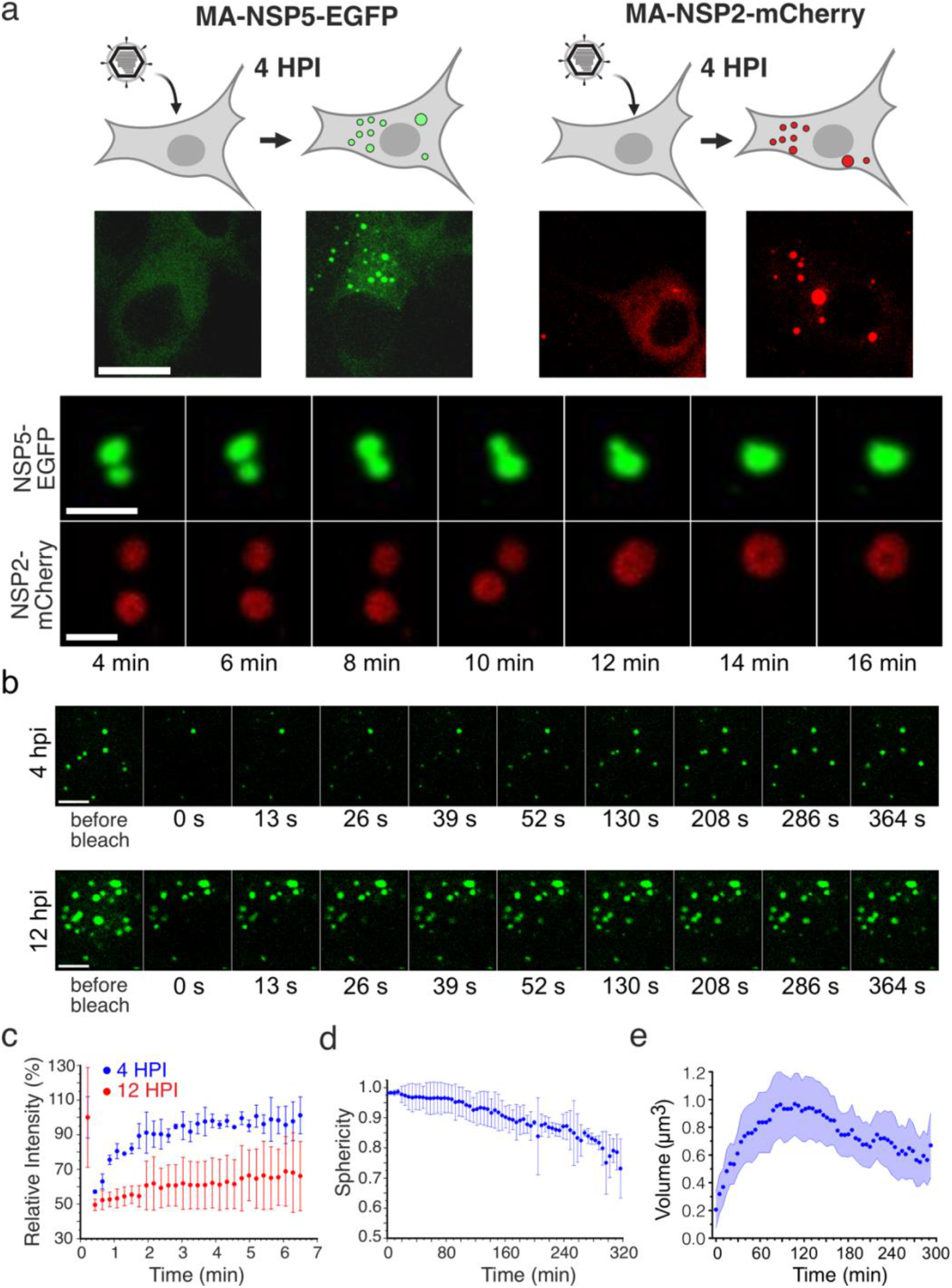
Liquid-like Properties of the Rotavirus Replication Factories **(a)** Dynamics of replication factories tagged with EGFP (NSP5-EGFP) and mCherry (NSP2-mCherry) visualised in MA104-NSP5-EGFP and MA104-NSP2-mCherry rotavirus-infected cell lines. Live-cell confocal images (4-16 min) acquired after 4 hours post infection (HPI). Scale bars, 5 μm. **(b)** Fluorescence recovery after photobleaching (FRAP) of EGFP-tagged replication factories after 4 HPI (early infection) and 12 HPI (late infection). **(c)** Fluorescence Intensities after FRAP of EGFP-tagged replication factories after 4 HPI (blue) and 12 HPI (red) shown in **(b)**. Each data point represents mean±SD intensity values calculated for multiple NSP5-EGFP-tagged granules in 5 RV-infected cells. **(d)** Sphericity of NSP5-EGFP-containing granules during RV infection. Each data point represents mean±SD sphericity values calculated for NSP5-EGFP-NSP5-tagged granules in cells detected in 15 frames. Data were recorded for 320 min immediately after 4 HPI when multiple NSP5-EGFP granules could be detected in RV-infected cells. **(e)** Calculated volumes of NSP5-EGFP-tagged granules formed in RV-infected cells after 4 HPI as shown in **(d)**. The mean values decrease due to *de novo* formation of multiple smaller NSP5-EGFP granules that continuously assemble in cells between 4 HPI (t=0 min) and 9 HPI (t=300 min).

To assess the dynamics of NSP5-EGFP in these droplets, we photobleached viroplasms during ‘early’ (4 HPI) and ‘late’ (12 HPI) infection and measured fluorescence recovery over time (**Fig. 1b**). Fluorescence recovery after photobleaching (FRAP) studies of the ‘early’ viroplasms revealed a rapid (60–80 s) and complete (95–100%) fluorescence recovery. The kinetics and recovery percentage, however, decreased substantially for larger granules observed during late infection stages (**Fig. 1c**). The reduction in FRAP recovery suggests a change in the viscoelasticity of late viroplasms (i.e., they become more viscous or solid-like) during the course of infection.

As most typical properties of liquids are determined by their surface tension^24,25^, smaller liquid droplets coalesce and attain spherical shapes with the lowest volume-to-surface area ratios. To investigate the shape of viroplasms, we observed NSP5-EGFP-expressing RV-infected cells, and we found that at 4 hours post infection these structures are spherical (**Fig. 1d** and Materials and Methods). Time-resolved confocal microscopy of individual viroplasms (Materials and Methods) revealed that size of droplets increased over course of infection (**Fig. 1a**). In contrast, the calculated sphericities of these inclusions decreased with time, suggesting loss of fluidity, consistent with the observed slower FRAP recovery rates during late infection (**Fig. 1c**).

We next examined the sensitivities of both early and late viroplasms towards the aliphatic alcohol 1,6-hexanediol (1,6-HD) which is commonly used as a chemical probe to differentiate between liquid-like and gel-like states of membrane-less organelles^26,27^. We exposed cells infected with rotaviruses to 4% (*w*/*v*) 1,6-HD added to cell culture medium. Immediately after application of the compound (less than 30 s), early viroplasms were completely dissolved (**Fig. 2a**). When 1,6-HD was removed, NSP5-EGFP assemblies slowly reappeared, initially forming smaller assemblies that eventually coalesced into larger viroplasms (**Fig. 2a** and Supplementary Movie 1). In contrast, when treated with 1,6-HD at 12 HPI, only a fraction of smaller viroplasms were dissolved, while larger viroplasms remained unaffected (**Fig. 2a**), suggesting that they have undergone a liquid-to-solid phase transition^28^. A brief (5 min) chemical crosslinking with 4% (*v*/*v*) paraformaldehyde prior to the application of the aliphatic alcohol also rendered the early infection (4 HPI) structures refractory to 1,6-HD treatment (Supplementary Figure 1a). Collectively, these results suggest that the assembly of viroplasms is driven by weak hydrophobic interactions that can be stabilised by chemical cross-linking. Additionally, we verified the 1,6-HD sensitivity of viroplasms assembled in the RV-infected cells producing NSP2-mCherry in lieu of NSP5-EGFP (Supplementary Figure 1b and Supplementary Movies 2 and 3). Irrespective of the protein tagged (NSP5 or NSP2), or the fluorophore chosen, viroplasms responded similarly to the application of 1,6-HD.

**Figure 2.**
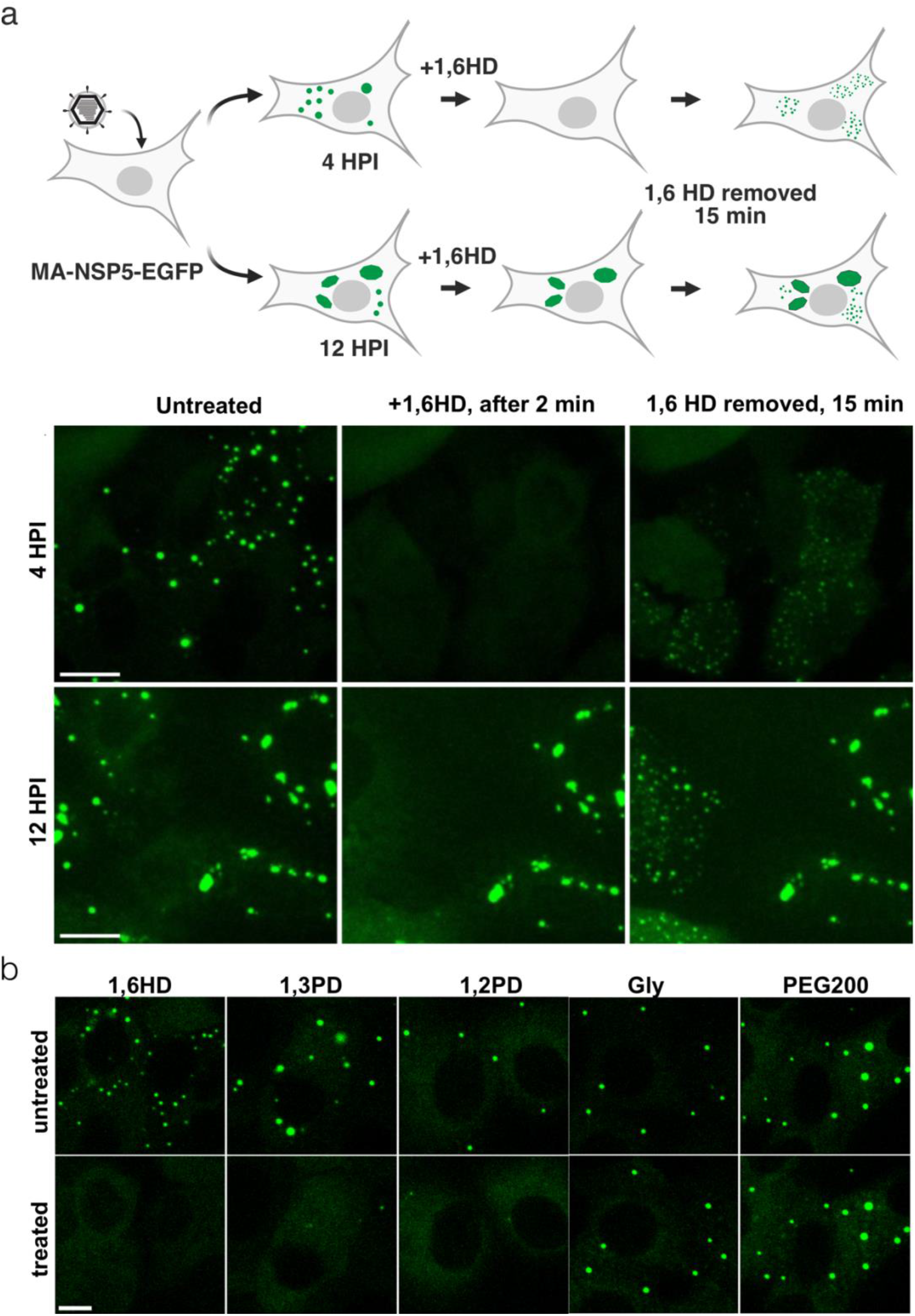
1,6-hexanediol (1,6HD) differentiates early and late viral replication factories **(a)** Replication factories in MA104-NSP5-EGFP cells infected with RV at 4 HPI dissolve after >30 s post application of 4% (w/v) 1,6HD added to the cell culture medium (middle panel). Removal of 1,6HD results reassembly of multiple EGFP-NSP5-containing droplets dispersed in the cytosol (right panel). Bottom, replication factories at 12 HPI: before application of 1,6HD (left), 2 min after application (middle), and 15 min after removal of 1,6HD from cell culture medium (left). Note larger viral factories that remain refractory to 1,6HD treatment. Scale bar, 50 μm. **(b)** Sensitivity of RV replication factories to aliphatic alcohols at 4 HPI. Left to right –1,6-hexanediol (1,6HD); 1,3-propylene diol (1,3PD); 1,2-propylene diol (1,2PD, or propylene glycol); glycerol (Gly); polyethylene glycol 200 (PEG200). Top panels – before application and bottom panels – 1 min after application of these compounds (4% v/v). Scale bar, 30 μm.

### Rotavirus Replication Factories Are Dissolved by Small Aliphatic Diols

We posited that related aliphatic diols with similar physicochemical properties (e.g., hydrophobicity and molecular weight) to 1,6-HD, but less toxic for cells might exert similar effects on these condensates in cells. Remarkably, we identified two low molecular weight aliphatic diols (1,2- and 1,3-propane diols; denoted as 1,2-PD and 1,3-PD, respectively) that also dissolved viroplasms in RV-infected cells at 4 HPI (**Fig. 2b**).

Since both intracellular protein concentration and protein tagging may significantly affect the properties of the phase-separating system^10^, we also carried out immunofluorescent staining of wild type MA104 cells infected with wild type RV before and after application of 1,6-HD and a non-toxic 1,2-PD (commonly known as propylene glycol, PG, Supplementary Figure 1c). Both alcohols completely dissolved viroplasms, further corroborating that the observed structures are formed via LLPS of NSP5 that accumulates during rotavirus infection.

As a final test, we used a recombinant NSP5-defficient (knock-out, KO) rotavirus^16^ to infect three MA104 cell lines that stably produce NSP5, NSP5-EGFP and NSP2-mCherry. Viroplasms were only observed in the cells producing untagged NSP5 4-8 HPI (Supplementary Figure 1). In contrast, no viroplasms were detected in NSP2-mCherry and NSP5-EGFP cells, confirming that the untagged NSP5 is the key protein that drives LLPS. Together with our recent studies^16^, these results also suggest that C-terminal tagging of NSP5 impairs its function and RV replication, whilst not precluding NSP5-EGFP mixing with untagged NSP5/NSP2 condensates that are formed during RV infection. Viroplasms sensitive to 1,2-PD treatment were also formed in mouse embryonic fibroblasts several hours post RV infection (Supplementary Figure 2), suggesting that the observed NSP5-rich condensates are formed via LLPS in other cell types susceptible to RV infection.

Taken together, early infection stage viroplasms exhibit all the hallmarks of a liquid state: they are spherical and they coalesce; they exchange cytoplasmically dissolved proteins; they are reversibly dissolved by the aliphatic alcohols disrupting weak interactions that drive LLPS.

### Rotavirus Viroplasm-forming Nonstructural Proteins Form Liquid Condensates

To move towards a better mechanistic understanding of liquid–liquid demixing of the two viroplasm-enriched proteins NSP5 and NSP2, we analysed their propensities to undergo LLPS. The high content of intrinsic disorder of NSP5, and its Gly/Ser and Asn/Glu-rich composition contribute to a number of several low complexity regions that typically favour weakly self-adhesive interactions required for phase separation. To this end, we first analysed the two proteins using our recently developed machine learning approach termed DeePhase^29^ to identify LLPS-prone sequences. This hypothesis-free approach revealed several protein regions (LLPS score > 0.5) with high propensity for driving phase separation (**Fig. 3a** and **b**). Remarkably, sequences of NSP5, predicted to promote LLPS, were located within the two regions of the protein that had been previously demonstrated to be essential for viroplasm formation^17^ (**Fig. 3a**, regions highlighted in green). One of these regions contained multiple negatively charged residues (**Fig. 3a,** C-terminal negatively charged residues shown in red), previously proposed to interact with the positively charged surfaces of NSP2^30^. Similarly, DeePhase detected multiple NSP2 residues predicted to have high propensity to drive LLPS (**Fig. 3b**). Remarkably, structural analysis^31^ of the octameric RNA chaperone NSP2 revealed that the majority of these residues presented multiple positively charged side chains (**Fig. 3c**), including those previously demonstrated to bind NSP5^30^. Given the highly charged natures of both proteins (NSP2, pI ~9 and NSP5, pI ~5.5), the observed phase separation is likely to be driven by both attractive electrostatic interactions (**Fig. 3c**), as well as hydrophobic interactions that are perturbed in the presence of the aliphatic alcohols, e.g., 1,6-HD and PG.

**Figure 3.**
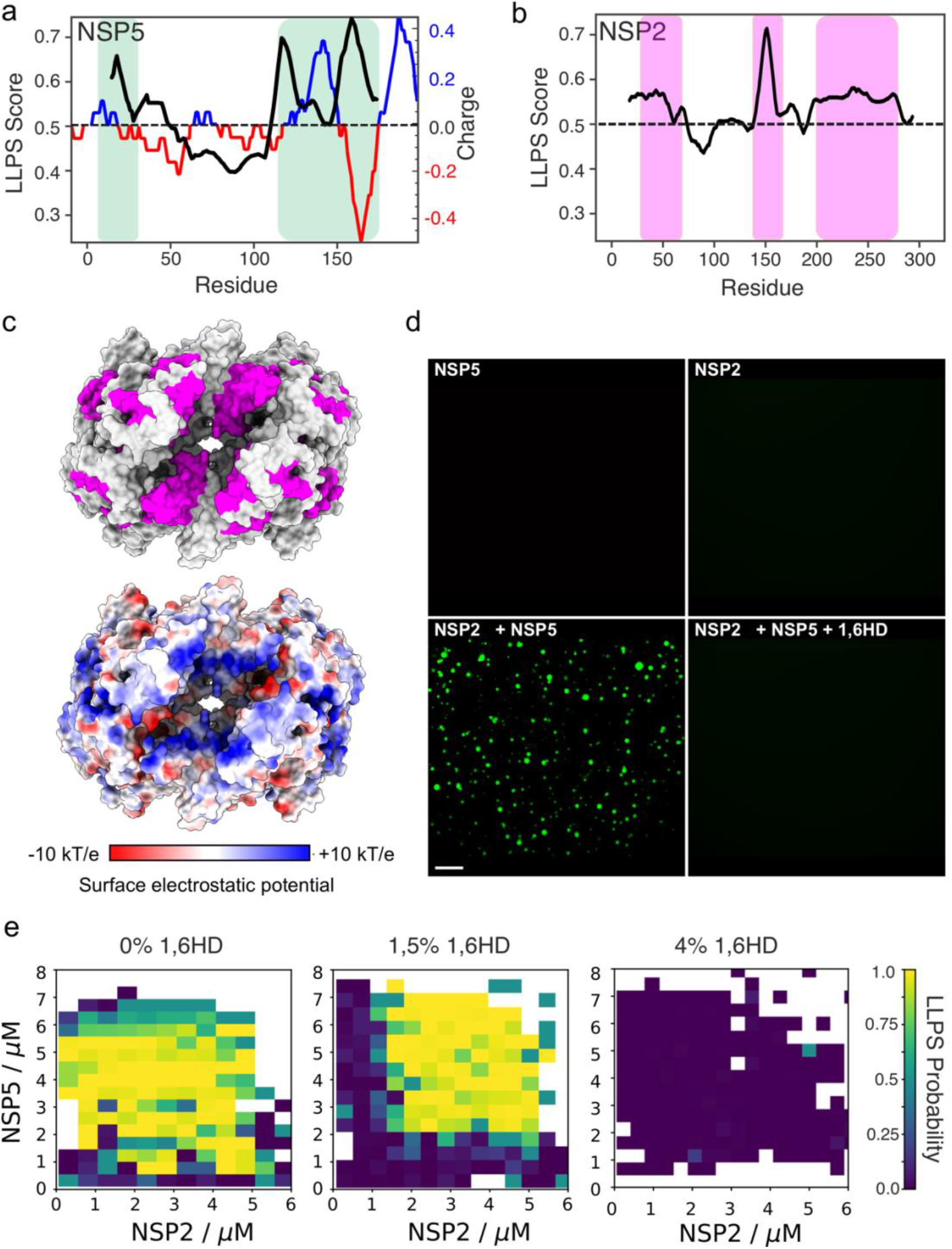
Rotavirus proteins NSP5 and NSP2 undergo liquid-liquid demixing. **(a)** DeePhase prediction of the phase-separating properties of NSP5 (LLPS score of 0.6). Averaged score with a 30-residue sliding window is shown as a black line. EMBOSS Protein charge plot shown as an alternating blue (positive charge) and red (negative charge) line. Green boxes denote protein residues of previously shown to be essential for viral factory formation in RVs. **(b)** DeePhase prediction of the phase-separating properties of NSP2 (LLPS score of 0.3). Note residues with high propensity to undergo LLPS (highlighted). These NSP2 residues were mapped on the surface representation of the RNA chaperone NSP2 in magenta **(c,** *top*) and for comparison shown along with the surface electrostatic charge distribution of NSP2 (**c**, *bottom*). **(d)** Atto488-labelled NSP5 (*top left*) and NSP2 (*top right*), 10 μM each, and immediately after mixing (*bottom left*). NSP5/NSP2 droplets are dissolved in the presence of 5% 1,6HD (*bottom right*). Scale bar, 20 μm. **(e)** Phase diagrams generated through droplet microfluidics for the coacervation of NSP2 and NSP5, in the presence of 0% v/v (*left*), 1.5% v/v (*middle*), and 4% v/v (*right*) 1,6-hexanediol. Phase diagrams were generated from N = 2206, 2035 and 1470 data points for each 1,6-hexanediol concentrations, respectively, and the data were used to construct the LLPS probability plots.

To confirm these results from our *in-silico* predictions, we next examined the behavior of the recombinant untagged NSP5 and the C-terminally His-tagged NSP2. In solution, each protein was monodisperse (Materials and Methods). Circular dichroism analysis of NSP5 suggested that regions of protein disorder contributed to almost 40% of the spectrum (Supplementary Figure 3). Such low-complexity intrinsically disordered regions commonly underpin LLPS of proteins^9,24,32,33^, in agreement with *in silico* predictions.

In isolation, NSP5 and NSP2 samples did not form any microscopically detectable condensates in the micromolar concentration regime (**Fig. 3d**, upper panel). Immediately upon mixing, 5–10 μM of each protein, multiple micron-sized droplets were formed (**Fig. 3e**). These droplets were dissolved by 1,6-HD (**Fig. 3e**), confirming that the observed droplets represented NSP5/NSP2 condensates, consistent with the effects of the aliphatic alcohol *in vivo.*

To further characterise the phase behavior of NSP5/NSP2 condensates, we generated phase diagrams for these protein mixtures alone and in the presence of 1,6-HD. Using high-throughput droplet microfluidics (Supplementary Figure 4), we obtained phase diagrams for a range of NSP5 and NSP2 concentrations (**Fig. 3e** and Supplementary Figure 5), revealing coacervation of the proteins occurred in the low micromolar regime. NSP5/NSP2 protein mixtures remained homogenous in the presence of 4% (*w*/*v*) 1,6-HD, with a detectable change in the phase-separation behavior observed even at lower 1,5% (*w*/*v*) 1,6-HD concentration. Given the potential electrostatic contribution of negatively and positively charged residues of NSP5 and NSP2, we also examined the salt-dependence of the NSP5/NSP2 condensate formation *in vitro.* Above 0.5 M NaCl concentration, NSP5/NSP2 coacervation was severely inhibited (Supplementary Figure 5), supporting the idea that NSP5/NSP2 coacervation is driven by both hydrophobic and electrostatic interactions between the two multivalent proteins.

### Viroplasms Are Complex Ribonucleoprotein Condensates That Accumulate RV RNAs

Given that viroplasms are viewed as sites of viral RNA replication^13,34^, and NSP2 is a multivalent RNA chaperone^35–37^, we then examined RNA composition and the effects of aliphatic alcohols on RNA distribution in viroplasms. smFISH analysis of the RV genomic segment 3 (Seg3) and segment 4 (Seg4) transcript distribution in MA104-NSP5-EGFP cells revealed that NSP5-EGFP marked viroplasms contained both types of transcripts at 4 HPI (**Fig. 4a,** *left*). Treatment of RV-infected cells with an aliphatic diol PG resulted in rapid disassembly of these ribonucleoprotein granules and re-localisation of the RV transcripts into the cytoplasm (**Fig.4a**, *middle*). Removal of these compounds from the cell culture medium for 15 min prior to cell fixation and imaging permitted reformation of smaller NSP5-EGFP granules containing both Seg3 and Seg4 transcripts. Confocal microscopy of the RNA signals suggests that Seg3 and Seg4 RNA transcripts remained intact upon viroplasm dissociation, consistent with rapid (~15 min) reformation of multiple RNA-rich granules upon removal of PG.

**Figure 4.**
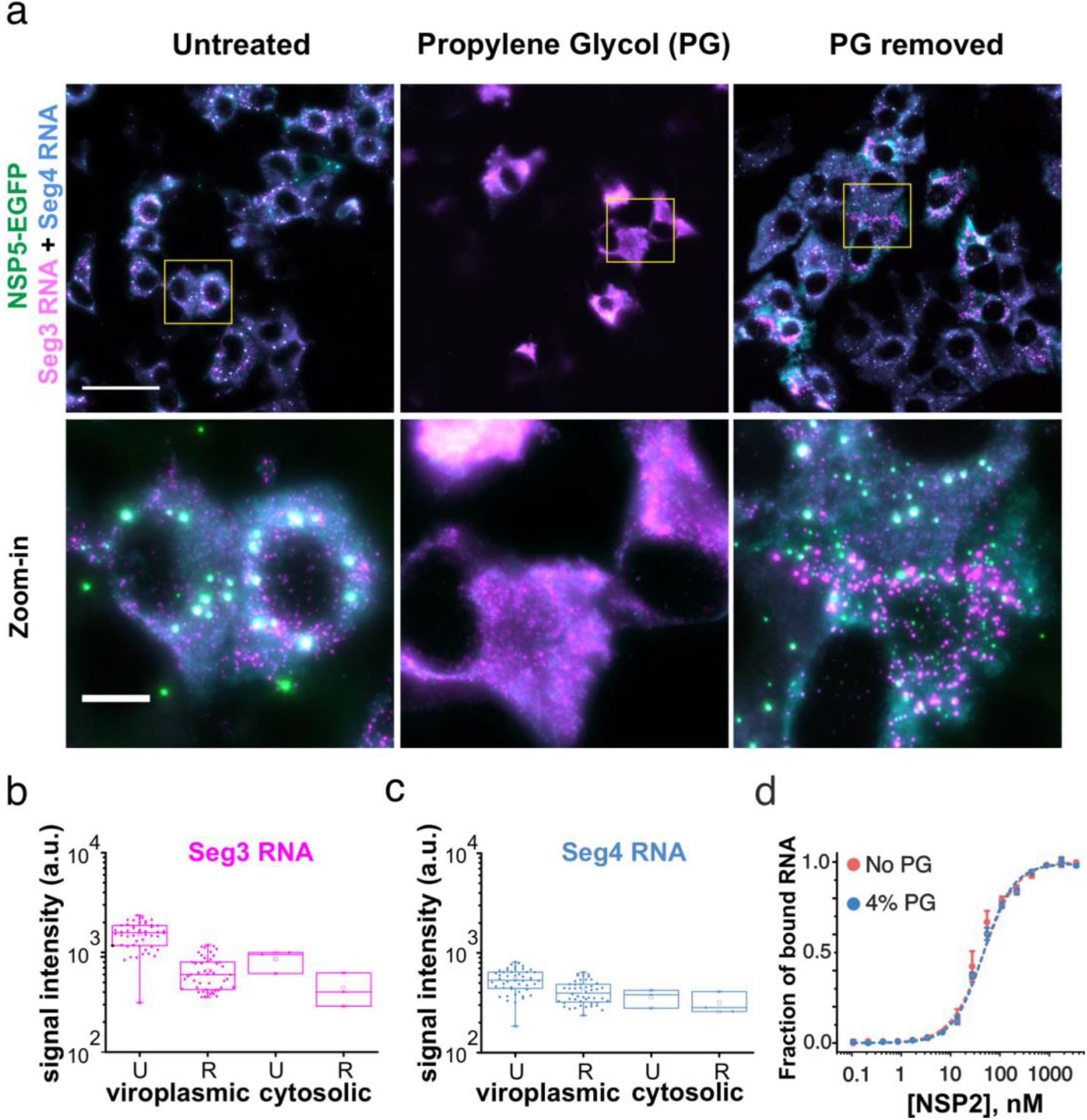
Rotavirus replication factories are RNA-protein condensates sensitive to propylene glycol. **(a)** RV-infected MA-NSP5-EGFP cells at 6 HPI. NSP5-EGFP-tagged viral factories (green) are rapidly dissolved in the presence of 4% (v/v) propylene glycol (PG, *middle*). Viral RNA-protein condensates rapidly reform (<10 min) after removal of propylene glycol (PG) from the cell culture medium (*right*). Rotavirus transcripts Seg3 RNA (magenta) and Seg4 RNA (cyan), with with colocalising Seg3 and Seg4 RNA signals (white) were detected by smFISH. Scale bars: 50 μm, zoomed-in regions: 10 μm. **(b-c)** Changes in the localisation of Seg3 and Seg4 RNAs and their relative distribution between the viroplasms and the cytosol before (Untreated, U), and 15 min after PG treatment (Recovery, R). Median and quartile values of integrated signal intensities (normalised by area) for each channel for viroplasms (‘viroplasmic’), and individual cells (N=9, ‘cytosolic’) are shown. **(d)** Binding of NSP2 to a fluorescently labelled 20-mer ssRNA in the presence of 4% propylene glycol (PG), measured by fluorescence anisotropy.

A fraction of RV transcripts formed multiple RNA clusters outside NSP5-EGFP granules (**Fig. 4a)**, suggesting that the viral transcripts aggregate independently of the ability of NSP5 and NSP2 to form liquid condensates. Analysis of the integrated RNA signal intensities before and after PG treatment revealed that the amount of transcripts did not dramatically change during these treatments (**Fig. 4 b and c**), confirming their reversible redistribution in the cytoplasm of infected cells. Interestingly, after recovery not all NSP5/NSP2 condensates were equally enriched in viral RNAs, further corroborating viral RNA re-distribution and exchange between these granules. Our recent studies indicate that rotavirus RNA oligomerisation is dependent on NSP2^35,38^.

The apparent affinity of NSP2 for RNA was identical in the presence of 4% (*v*/*v*) PG (**Fig. 4d**), suggesting that despite the observed perturbation of the NSP5/NSP2 condensates with aliphatic diols, NSP2–RNA complexes did not dissociate under those conditions.

This aspect of viroplasm formation remarkably resembles the formation of other complex ribonucleoprotein condensates, e.g., paraspeckles, in which RNA foci did not dissociate in the presence of aliphatic diols, despite the apparent dissolution of paraspeckles^39^. We therefore characterised the RNA foci formed in RV-infected cells during early infection using super-resolution DNA-PAINT approach^40^ combined with smFISH. This super-resolution technique exploits transient binding of fluorescent DNA probes (‘imagers’) to complementary, RNA-bound ‘docking’ DNA strands (**Fig. 5a** and **b**). At 4 HPI, Seg3 transcripts could be detected as submicron-sized RNA clusters (**Fig. 5d**), similar to those seen in diffraction-limited images (**Fig.4a**). 3D DNA-PAINT imaging of NSP2 condensates in RV-infected cells confirmed that early infection condensates contain only few viral transcripts, suggesting that NSP5/NSP2 coacervation spontaneously occurs during early RV infection, and it is not nucleated by the transcribing viral particles present in cells. Furthermore, 3D DNA-PAINT imaging of Seg3 RNA foci revealed that they became less isotropic (i.e., loss of sphericity) by 6 HPI (Supplementary Figure 6), mirroring the overall decrease in sphericity of viroplasms during late infection. Given the resistance of RNA foci to aliphatic diols, and rapid (10–15 min) reformation of smaller condensates upon removal of these compounds, it is possible that such viral RNA aggregates could seed the nucleation of new NSP2/NSP5 condensates in cells^41^. Taken together, these results confirm that early infection viroplasms should be regarded as liquid RNP granules^42–45^.

**Figure 5.**
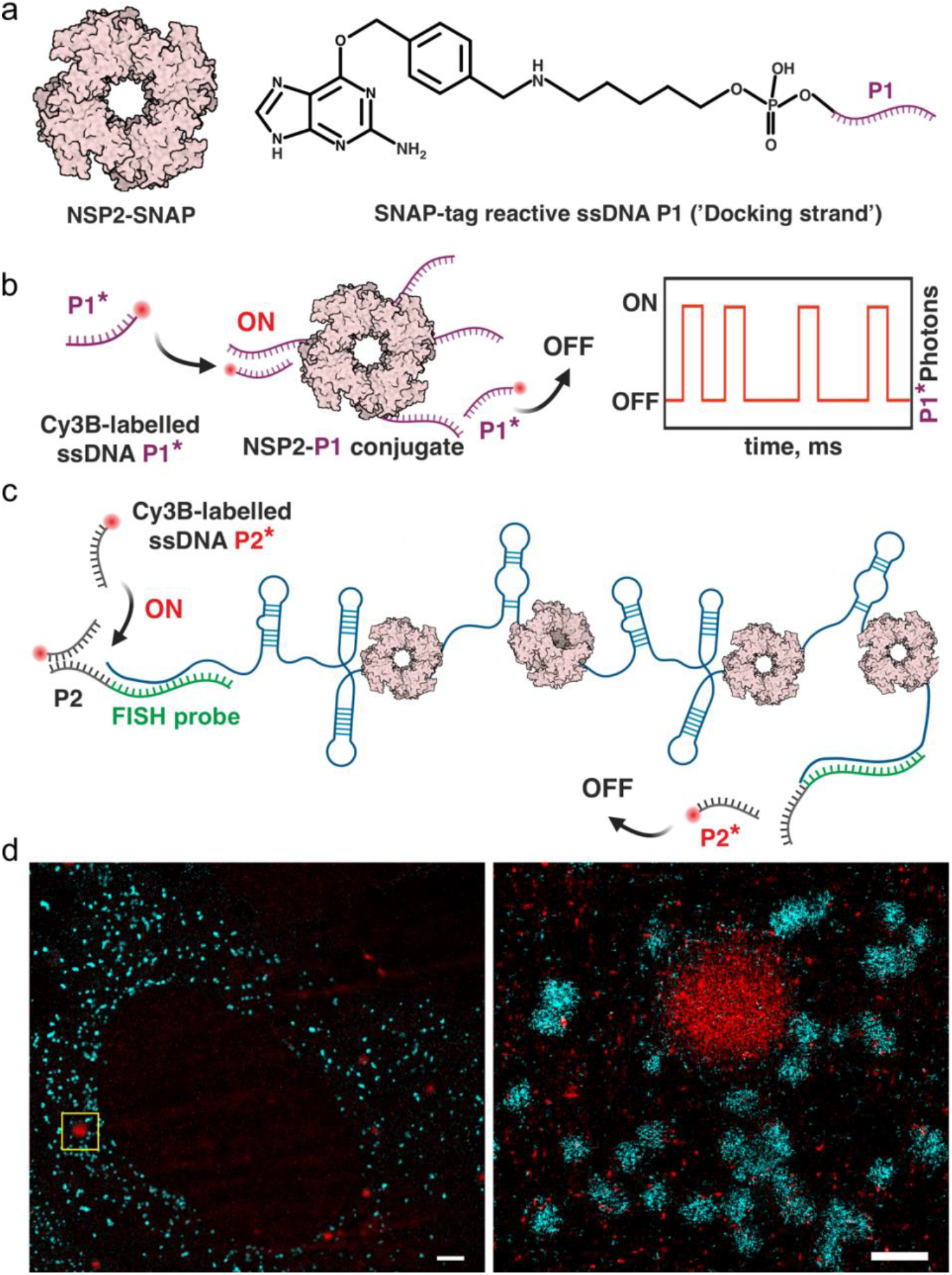
Super-resolution DNA-PAINT analysis of RV replication factories. **(a)** DNA-labelling scheme of NSP2 for DNA-PAINT imaging. Low levels of SNAP-tagged NSP2 (a doughnut-shaped NSP2 octamer) are produced in a stable MA104 cell line. A SNAP-tag reactive benzylguanine (BG) DNA derivative (P1 ssDNA docking strand) can form a stable thioether bond with NSP2-SNAP. **(b)** Detection of NSP2-rich condensates formed in RV-infected cells using DNA-PAINT approach. Transient binding and dissociation of a Cy3B-dye labelled ssDNA probe P1* (complementary to ssDNA P1) generates blinking at the target sites (ON/OFF) used for stochastic super-resolution imaging. **(c)** Similarly, smFISH of Seg3 transcripts is combined with DNA-PAINT approach by installing orthogonal P2 ssDNA docking sites into the Seg3-specific FISH probes **(d)** A combined super-resolved image of NSP2-rich condensates (NSP2-SNAP in red) and Seg3 transcripts (cyan) in RV-infected cells 4 HPI. Scale bars = 2 μm (*left*), 500 nm (*right*).

## Discussion

Previous studies uncovered protein composition of viroplasms, suggesting that these cytoplasmic inclusions are formed when NSP5 is co-expressed with NSP2 and/or the viral capsid protein VP2^14,15,17,23,46^, even in the absence of RV infection. Here, from multiple lines of evidence, we revealed that rotavirus viroplasms represent ribonucleoprotein condensates that are formed via phase separation of NSP5 and NSP2: First, cytosolic inclusions formed by NSP5/NSP2 are initially spherical; second, two or more NSP5-rich droplets can fuse and relax into a sphere; third, infection with the NSP5-defficient rotavirus mutant does not yield these droplets, unless NSP5 is produced *in trans*; fourth, these droplets are formed upon NSP5 and NSP2 mixing *in vitro*, and when these proteins are co-expressed *in vivo;* finally, NSP2/NSP5 droplets can be instantly dissolved when treated with aliphatic alcohols known to disrupt multivalent interactions driving liquid–liquid phase separation. Our findings that distinct compounds other than 1,6-hexanediol but with similar physicochemical properties can reversibly dissolve viroplasms in RV-infected cells reinforces the idea that these inclusions represent liquid condensates. Furthermore, these compounds did not dissolve viroplasmic condensates after chemical cross-linking. Importantly, the observed condensates are formed exclusively when the untagged version of NSP5 is co-expressed in cells along with either NSP2, or VP2^17,46^, as well as during RV infection. Thus, the formation of the observed condensates reflects the unique physicochemical properties of NSP5/NSP2, and is not artifactual due to over-expression or fluorescent tagging^47,48^.

NSP5 and NSP2 have been long established to be major binding partners and the key protein residents of RV replication factories^12,14,16,17^. Given the multivalent RNA and NSP5-binding nature of NSP2, the observed phase-separation of these proteins at low micromolar concentration is consistent with the reports of their aggregation-prone behaviour upon mixing at higher micromolar concentrations^30,35^. By exploring the phase boundary using the PhaseScan, we have shown that the degree of NSP5/NSP2 coacervation is determined by the concentrations of both interacting partners. Thus, our model predicts that the kinetics of NSP5/NSP2 condensate formation depends on the intracellular concentration of both NSP5 and NSP2, the production of which would be expected to directly correlate with the number of infectious particles per cell. Indeed, previous observations^49^ suggesting that the kinetics of viroplasm formation increase in direct correlation with the multiplicity of infection fully support our model.

Current views of replication factory formation in RVs are dominated by the idea of multiple viral proteins being recruited into viroplasms in a specific order^15,17,23,46,50^ resulting in their particular organisation^51^. Here, we propose a unifying model for viroplasm assembly (**Fig. 6**) that takes into account extensive existing data on their structural organisation accrued over several decades, and amounts to a step change in our understanding of these replication factories in these viruses. We propose that RV viroplasms represent condensates formed by NSP5/NSP2 coacervation. Initially, these condensates behave as dynamic fluids, and change in their viscoelastic properties (e.g., fluidity) during infection, concomitant with changes in viral protein phosphorylation^15,16,23,52–55^ and the ratio of RNA:protein in these inclusions. Other cellular biomolecular condensates have been shown to contain hundreds of distinct molecular species^56^, acting as membraneless protein-rich liquid condensates that selectively partition biomolecules and can promote specific nucleic-acid remodeling events^6^. Despite their complex and dynamic composition, typically, only a few protein residents are required to form these condensates^9,10,56,57^. Given its high propensity to undergo LLPS, and multiple lines of evidence demonstrating its indispensable role in the formation of viroplasms^12,16,17,52,53^, we propose that the rotavirus NSP5 acts as the primary scaffold required to form these condensates. Knocking out NSP5 abolishes formation of these structures even when other viral proteins are present during infection^16^ (Supplementary Figure 1), while NSP5 co-expression with RV multivalent RNA-binding proteins, e.g., NSP2, ^17,21^ results in formation of such condensates. Our *in vitro* results fully corroborate the model for NSP5/NSP2 condensate formation, further suggesting that the observed condensation does not require phosphorylation of proteins *in vitro*, despite multiple phosphorylated forms of NSP5 detected in rotavirus-infected cells^16,52,54^. This result suggests the phosphorylation of NSP5 is not a prerequisite for coacervation, but it is likely to play important roles in regulation of the molecular selectivity and specificity of these condensates, thus allowing their protein and RNA composition change during infection. Similarly, phosphoregulation of protein condensates has been demonstrated for a number of membraneless organelles *in cellulo* and *in vitro*^58,59^. Other condensate residents are concentrated within these coacervates, often by direct interactions with scaffolds, but are not required for condensate formation and referred as clients^56^.

**Figure 6.**
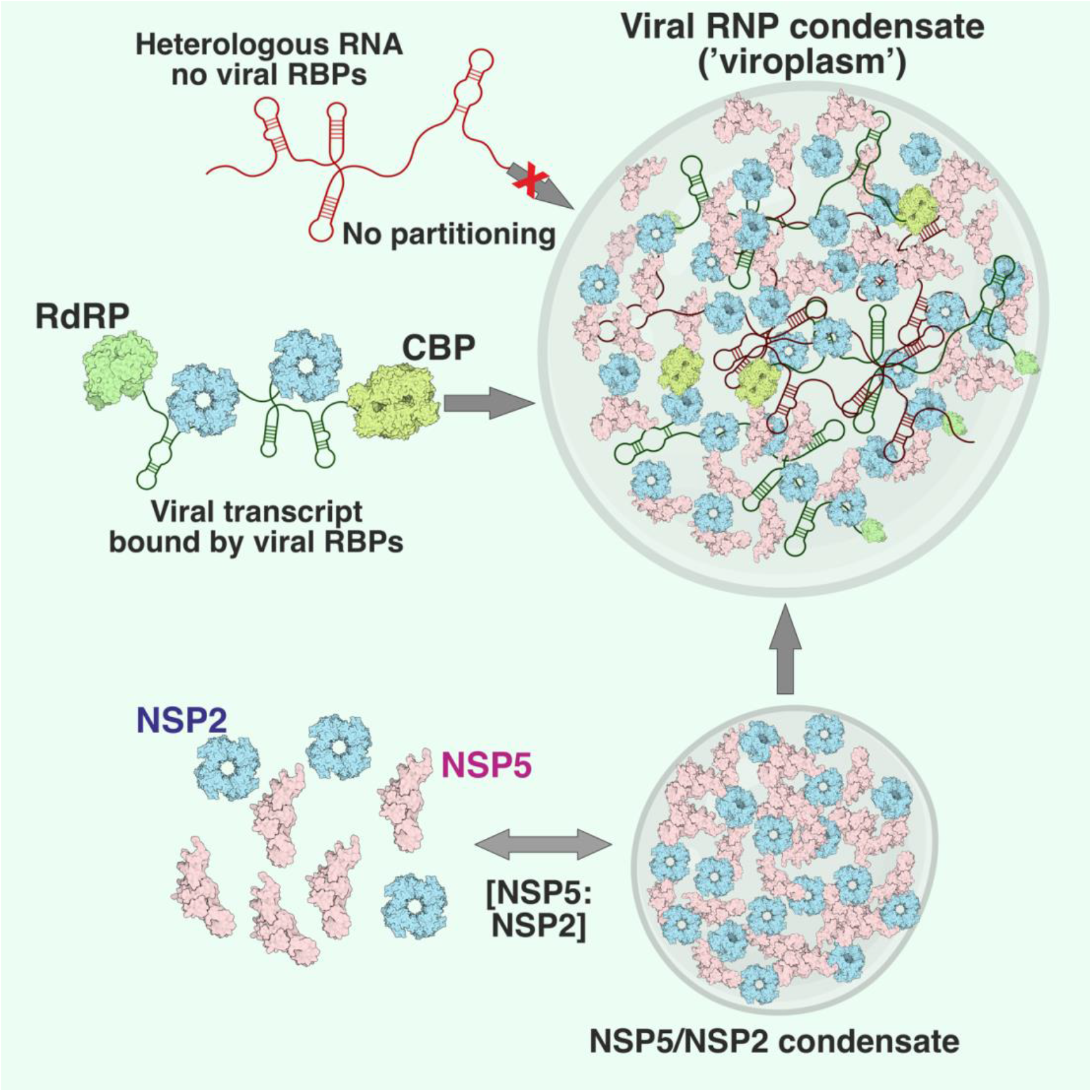
Proposed model of LLPS-driven formation of viral replication factories in rotaviruses. Multivalent Asp/Glu- and Ser-rich protein NSP5 (pink) acts as a scaffold that binds multivalent RNA chaperone NSP2 (cyan doughnut-shaped octamers), and potentially other RNA-binding clients. NSP5 and NSP2 readily undergo coacervation at low micromolar concentrations, also forming condensates in cells previously described as ‘viroplasm-like structures’. Cognate viral transcripts undergo enrichment in these condensates via a mechanism distinct from other known RNP granules. Mechanistically, this could be achieved by specific protein-RNA recognition, e.g., the viral RNA-dependent RNA Polymerase (RdRP) that recognises conserved sequences of all eleven distinct RV transcripts, and undergoes partitioning into NSP5/NSP2 condensates due to its high nM affinity for both proteins^1,2^. Other similarly sized RNAs (red) lacking the viral proteins with high affinity for NSP5/NSP2 proteins are excluded from these condensates. Other multivalent RNA-binding proteins (RBPs), i.e., the viral cap-binding protein (CBP), and multiple copies of NSP2 can form the RNP complexes that can be absorbed into the NSP2/NSP5 condensates, and released from them by dissolving the condensates. Such RNP condensates are known to promote RNA-RNA interactions in cells, while simultaneously acting as ‘molecular filters’ for concentrating viral transcripts, both mechanisms being conducive to a multi-RNA genome assembly in rotaviruses.

Viral RNA-binding proteins (e.g., viral RNA-dependent RNA polymerase, RdRP, and a cap-binding protein, **Fig. 6**) concentrate in these structures, and have been demonstrated to strongly bind NSP5^50,60^, but not being sufficient to form viroplasm-like structures on their own, thus fulfilling the criteria of client proteins that partition into NSP5/NSP2 condensates. Lipid bilayers^61^, microtubules^62^ and tubulin^63^ can promote nucleation of biomolecular condensates and spatially regulate the kinetics of their formation in cells. Association of lipid droplets^64,65^ and tubulin^15,49^ with viroplasms is thus entirely consistent with our model of formation of viral replicative factories in RV-infected cells. Similarly to the early and late infection stage viroplasms, changes in the viscoelastic properties of liquid condensates that decrease fluidity over time are common for many other systems that undergo LLPS^28,66,67^. Recent super-resolution imaging studies of these organelles in RV-infected cells proposed that distinct viral proteins are organized into multiple concentric layers^51^. The proposed model explains the relevance of these findings, as even very simple condensates show characteristics of multilayered behavior^33^. Distinct layers are likely to form via different molecular interaction networks that lead to different viscoelastic properties, such as those observed in nucleoli^68^, P-granules^32^ and nuclear speckles^69^.

### Implications for selective RNA recruitment and RNA-RNA interactions required for segmented genome assembly

Coacervation of the viral RNA chaperone NSP2^35,38,70^ associated with multiple viral transcripts would accelerate formation of inter-molecular RNA–RNA interactions between them. Several processes may contribute to stabilisation of sequence-specific inter-molecular RNA-RNA contacts^44,71^, while promoting intra-molecular duplex melting via interactions with multiple arginine side chains of NSP2^72^ concentrated in the viroplasmic liquid phase^6^. Recent evidence argues that intermolecular RNA-RNA interactions play a role in forming and determining the composition of distinct cytoplasmic, RNA-rich ribonucleoprotein granules^42–44,73,74^. Coalescence of multiple RNA-binding proteins and non-translating mRNAs lacking fixed stoichiometry can occur during cellular stress, giving rise to stress granules^73^. Similarly, viroplasms accumulate non-polyadenylated, untranslated viral transcripts and viral RNA-binding proteins. While stress granules are highly enriched in poly(A)-binding proteins associated with mRNAs, non-polyadenylated viral transcripts are likely to be bound by the viral RNA-dependent RNA polymerase (RdRP), previously reported to have nM affinity for both NSP2 and NSP5 (**Fig.6**). We propose that RV replication factories represent a unique case of specialized RNP granules that promote accumulation of viral transcripts to minimise spurious promiscuous RNA–RNA interactions with non-cognate host cell RNAs^44^. Our findings have several important ramifications for future studies of rotavirus replication mechanisms, posing many outstanding questions that arise from recognition of the LLPS-driven assembly of viroplasms, and potentially of other viral replication factories in segmented dsRNA viruses that exhibit similar liquid-like behavior in cells^75,76^. The proposed LLPS-driven mechanism of viroplasm formation offers a unified model that explains multiple results from previous efforts to explain their assembly, and establishes LLPS as an attractive target for antiviral intervention^77^.

## Materials and Methods

### Cells and Viruses

Rotavirus A strains (Bovine rotavirus strain RF and simian rotavirus SA11) were propagated as previously described^78,79^. MA104 (ATCC CRL-2378.1) and its derivatives MA-NSP2-mCherry and MA-NSP5-EGFP stable cell lines were generated and maintained as described in^16,17^. Lentiviral vector pAIP-NSP2-SNAP was generated using a synthetic SNAP tag-coding DNA (GenPart, Genscript) inserted into a double digested with *Mlu*I/*EcoR*I pAIP-NSP2-mCherry vector^16^. MA104-NSP2-SNAP cell line was then generated as previously described^16^. Briefly, 7×10^6^ HEK293T cells were seeded in 10-cm^2^ tissue culture dishes 24 h before transfection. For each well, 2.4 μg of pMD2-VSV-G, 4 μg of pMDLg pRRE, 1.8 μg of pRSV-Rev, and 1.5 μg of pAIP-NSP2-SNAP DNA constructs were co-transfected using Lipofectamine 3000 (Sigma-Aldrich) following the manufacturer’s instructions. After 48 h, the virus was harvested, filtered through a 0.45 mm polyvinylidene fluoride filter, and immediately stored at −80°C. For lentiviral transduction, MA104 cells were transduced in six-well plates with 1.2 ml of the lentivirus-containing supernatant for 2 days. Cells were then selected by growing cells in DMEM supplemented with 10% FBS and puromycin (5 μg/mL) for 4 days. NSP5 immunostaining was carried out as previously described^16^.

### Image data acquisition

Confocal imaging was conducted on a Zeiss Cell Observer SD inverted confocal microscope with a Yokogawa CSU-X1 spinning disk unit from Zeiss (Jena, Germany). The microscope was equipped with a 1.40 NA 63x Plan apochromat oil immersion objective from Zeiss. Measurements were performed at room temperature. Photo bleaching for FRAP experiments was done with a 488 nm laser at 100% intensity and 3000 ms exposure time, then the recovery was observed for 60 frames every 30 seconds. EGFP was imaged using a 488 nm laser at 20% intensity and 200 ms exposure time and mCherry was imaged with a 561 nm laser at 20% intensity and 200 ms exposure time. Images recorded as z-stacks consisted of either 10 or 50 frames, with a 0.5 μm distance between them, depending on the sample. In the excitation path a quad-edge dichroic beamsplitter (FF410/504/582/669-Di01-25×36, Semrock) was used. For two color detection of EGFP and mCherry a dichroic mirror (660 nm, Semrock) and band-pass filters 525/50 and 690/60 (both Semrock) were used in the detection path. Separate images for each fluorescence channel were acquired using two separate electron multiplier charge coupled devices (EMCCD) cameras (PhotometricsEvolve™). Image acquisition was controlled using the Zeiss Zen (blue edition) 2011 Software (Zeiss). Widefield imaging was conducted with the Eclipse Ti-E inverted microscope from Nikon (Tokyo, Japan). The images were acquired with a 0.7 NA 60x S Plan Fluor ELWD oil immersion objective from Nikon. Measurements were performed at room temperature. A pE-4000 illumination system (CoolLED) was used as light source. DAPI was imaged using a 385 nm LED at 33% intensity and 55 ms exposure time. EGFP and ATTO 488 were imaged using a 470 nm LED at 41% intensity with 300 ms exposure time and 7% intensity with 55 ms exposure time respectively. mCherry was imaged with a 550 nm LED at 36% intensity and 300 ms exposure time. The light path was regulated with a Dapi/FITC/Cy3/Cy5 Quad HC Filter Set (Semrock). The images were acquired using a scientific complementary metal–oxide–semiconductor (sCMOS) camera (Andor Technology). Image acquisition was controlled using the NIS-Elements AR V.4.50 Software (Nikon).

### Image Data Processing

The recorded pictures were processed with ImageJ (v.1.52p)^80^. Data of the FRAP experiments was also analysed with ImageJ. Distinct visible granules were selected manually as ROIs before bleaching. The recovery curve over a time span of 13 minutes (corresponding to 60 frames) was calculated for each ROI. The displayed values are median intensities of five ROIs. Other parameters, including fusion events, velocity or sphericity, were analysed with Imaris (v 8.2.0, Bitplane, AG Zurich, Switzerland). The viral granules were marked as ROIs based on their size and high fluorescence intensity. Then the centre of image mass R of the detected fluorescence volume in each ROI is calculated with the voxel (camera pixel) intensity m_i_, the center of a voxel r_i_ and the sum of voxel intensities M.

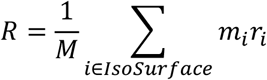

The coordinates of the centre of image mass of the ROIs was tracked for the duration of the experiment. From the resulting values the velocity was calculated as the change of the centre of image mass coordinates between two frames divided by the frame time (4.9 min). 65 ROIs were observed over a time span of 5.2 hours.

Sphericity Ψ was calculated as the ratio of the surface area of a sphere with the same volume as the given particle V_p_ to the surface area of the particle A_p_

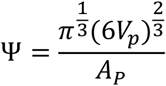

Mean sphericity values were calculated for 65 ROIs monitored for EGFP-marked granules observed for over 6 h. Fusion events were counted when two separate ROIs overlap and their volumes were treated as a single volume. 1037 ROIs were observed over 8 hours. If not stated otherwise, data points in the figures represent mean values averaged over all measured ROIs. Measurements were all performed in biological triplicates. The data presentation was done with OriginPro (Version 8.0891, OriginLab Corporation, Northampton, MA, USA). Where appropriate, schematics of figures were prepared using **BioRender.com**

### NSP5 and NSP2 expression and purification

Recombinant NSP2 (strains SA11 and RF) were expressed and purified, as previously described^35^. NTA-affinity purified NSP2 fractions were further purified over a HiTrap SP cation-exchange column. The concentrated peak fractions were resolved on a Superdex 200 10×300 GL column and pre-equilibrated with RNAse-free SEC buffer (25 mM HEPES-Na, pH 7.5, 150 mM NaCl) to ensure high purity and homogeneity of the preparation. While a functional form of the RNA chaperone NSP2 can be produced and purified under native conditions^31,35^, and its C-terminally His-tagged version supports viral replication^37^, previous attempts to natively purify a full-length untagged NSP5 were not successful^30,81^. We therefore expressed and purified NSP5 under denaturing conditions, followed by its refolding. Full-length recombinant NSP5 (strain RF) was expressed and isolated from bacterial pellets as inclusion bodies as previously described^35^. Washed inclusion bodies were solubilized in 6 M guanidinium hydrochloride and the protein-containing fraction was then subjected to a refolding protocol following step-wise dialysis^81^. After refolding, NSP5-containing fractions were further purified over an ImpRes Q column (GE). The concentrated peak fractions were further resolved on a Superdex 200 10×300 column pre-equilibrated with SEC buffer (25 mM HEPES-Na, pH 7.5, 150 mM NaCl) to ensure homogeneity of the preparation. Quasi-elastic scattering analysis of a monodisperse NSP5 sample revealed a hydrodynamic radius ~6.8 nm, consistent with the previously proposed decameric organisation^81^.

### DeePhase predictions

The propensity of the protein sequences to form condensates was estimated using the DeePhase model. Briefly, individual predictions relied on featuring the protein sequences by estimating a number of explicit sequence-specific parameters (sequence length, hydrophobicity, Shannon entropy, the fraction of polar, aromatic and positively charged residues and the fraction of sequence estimated to be part of the low complexity region and intrinsically disordered region) as well as implicit word2vec algorithm-based embeddings. The used model had been trained on previously constructed datasets including sequences with varying propensity to undergo LLPS as has been described in^29^. In order to evaluate how the LLPS-propensity of each protein sequence varied along its length, the full sequences were divided into 20 amino acid long fragments and the propensity of each fragment to undergo LLPS was evaluated. For the final result, individual predictions from 10 consecutive fragments were averaged.

### Circular Dichroism Spectroscopy and Dynamic Light Scattering

Samples were prepared by dialyzing NSP5 against 10 mM phosphate buffer pH 7.4, 50 mM sodium fluoride. Spectra were acquired in a 1 mm path length quart cuvette (Hellma) using a Chirascan plus spectrometer (Applied Photophysics) with a 1 nm bandwidth and a step size of 1 nm. An average of 3 scans (190-280 nm) were used for the final spectra, measured at 20ºC and 90ºC. Data were fitted to determine the secondary structure content using BeStSel^82^.

NSP5 samples (1 mg ml^−1^) were injected on a TSKgel G6000PWxl SEC column (Tosoh) pre-equilibrated with the SEC buffer (see above) at 21°C and a flow-rate set to 0.4 ml min^−1^. Dynamic (Quasi-Elastic) light scattering (QELS) measurements were carried out using an AKTA pure system (GE Healthcare) connected to a DAWN HELEOS and Optilab TrEX for QELS (Wyatt). On-line QELS was carried out using WyattQELS DLS Module to measure the translational diffusion and corresponding hydrodynamic radius of the eluting fraction. Autocorrelation functions (ACFs) were fitted to a single exponential to determine diffusion coefficients and corresponding hydrodynamic radii (Rh) of the oligomeric NSP5 species using ASTRA software (Wyatt).

### PhaseScan

#### Device fabrication

Polydimethylsiloxane (PDMS, Corning) devices for droplet generation and multilayer well-devices for droplet collection and imaging were produced on SU-8 (Microchem) moulds fabricated via photolithographic processes as described previously^83–85^.

#### Phase diagram generation

Phase diagrams were produced using droplet microfluidics in a similar manner to that described previously^19^. Syringe pumps (neMESYS modules, Cetoni) were used to control flows of protein solutions, consisting of 22 μM NSP5 supplemented with 6.4 μM Alexa647 dye (carboxylic acid, ThermoFisher) or 8 μM His-tagged NSP2 labelled with 8 μM Atto488-nitrilotriacetic acid (NTA, Sigma), and buffer (0.5 × phosphate saline buffer, PBS, pH 7.4). Appropriate quantities of 1,6-hexanediol were pre-mixed into all solutions before droplet generation. The aqueous flow rates were configured to vary automatically according to pre-set gradients, with constant total flow rate of 60 μL/h, to scan phase space between nominal concentrations of 0.9–7.3 μM and 0.30–6.5 μM for NSP5 and NSP2, respectively FC-40 oil (containing 1.5% (*v*/*v*) fluorosurfactant, RAN biotechnologies) was introduced to the device at a constant flow rate of 50 μL/h for microdroplet generation. For further details see Supporting Information.

#### Imaging

Directly after generation, microdroplets were transferred into a droplet-trapping device^86^ to ensure droplets were maintained in a well-spaced, stationary configuration for imaging. Microscopy data was acquired with a AxioObserver D1 microscope (Zeiss) equipped with a 5x air objective and a high-sensitivity camera (Evolve 512, Photometrics). Appropriate filter sets were used for EFGP (49002, Chroma Technology) and AlexaFluor 647 detection (49009, Chroma Technology). Representative data are presented in Supplementary Figure 5.

#### Droplet detection and data analysis

Acquired images were analysed using a custom-written Python script. Droplets were fitted as circles in the images. Non-circular droplets or erroneous detections were filtered and removed. From the fitted circular areas, the total intensity was calculated and normalised to obtain the intensity per unit volume (calculated using the fitted diameter), and converted to concentrations by comparison to calibration images acquired with known concentrations of NSP2/Atto488 and NSP5/Alexa647 mixtures. Droplets were classified as phase-separated or homogeneous according to the presence or absence of at least two connected pixels >5 standard deviations from the mean pixel intensity. Representative classification output is presented Supplementary Figure 5.Droplet classification, NSP2 and NSP5 concentration were then combined on a per-droplet basis to produce phase diagrams. Two-dimensional probability maps were constructed by division of the phase space (NSP2 vs. NSP5 concentration) into regular squares. The proportion of homogeneous or phase-separated droplets present in each region of phase space was calculated, before being passed through the error function (erf) to classify the phase-separation propensity of each region as represented by the colourmap.

### Affinity measurements by fluorescence anisotropy

Fluorescence anisotropy measurements with AlexaFluor488 dye-labelled 20-mer RNA, as described previously^70^, were performed at 25ºC using a POLARstar Omega plate reader (BMG Labtech) in Greiner 384 well black polypropylene plates. Serial 2-fold dilutions of NSP2 and σNS were titrated into 5 nM RNA in 50 mM Tris-HCl pH 7.5, 50 mM NaCl, 1 mM EDTA, 0.05% Tween-20 in a total volume of 50 μl and equilibrated at room temperature for 15 minutes prior to measurements were taken. Where required, buffers were supplemented with 4% v/v 1,2-propanediol. Raw Anisotropy (r) values were calculated as follows:

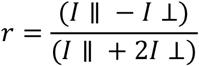

Where *I* ∥ and *I* ⊥ are the parallel and perpendicular emission signals, respectively. Normalized anisotropy values were plotted as a function of protein concentration and fitted to a Hill equation using OriginPro 9.0.

### Single-molecule Fluorescence in situ Hybridisation (smFISH) and TagPAINT

Rotavirus-infected and mock-infected MA104 cell controls, where appropriate, were fixed with 4% (v/v) methanol-free paraformaldehyde in nuclease-free phosphate saline buffer (PBS, Sigma) for 10 min at room temperature. Samples were then washed twice with PBS, and fixed cells were permeabilised with 70% (v/v) ethanol (200 proof) in RNAse-free water, and stored in ethanol at +4°C for at least 12 hours prior to hybridization, and no longer than 24 h. Permeabilized cells were then re-hydrated for 5 min in a pre-hybridization buffer (300 mM NaCl, 30 mM trisodium citrate, pH 7.0 in nuclease-free water, 10 % v/v Hi-Di formamide (Thermo Scientific), supplemented with 2 mM vanadyl ribonucleoside complex). Re-hydrated samples were hybridized with an equimolar mixture of DNA probes specific to the RNA targets (RVA strain SA11 and EGFP sequences), 62.5 nM final concentration, see SI Table 1, in a total volume of 200 μl of the hybridization buffer (Stellaris RNA FISH hybridization buffer, Biosearch Technologies, supplemented with 10% v/v Hi-Di formamide). After 4 hours of incubation at 37°C in a humidified chamber, samples were briefly rinsed with the wash buffer (300 mM NaCl, 30 mM trisodium citrate, pH 7.0, 10 % v/v formamide in nuclease-free water, after which a fresh aliquot of 0,3 ml of the wash buffer was applied to each well and incubated twice at 37°C for 30 min. After washes, nuclei were briefly stained with 300 nM 4’,6-diamidino-2-phenylindole (DAPI) solution in 300 mM NaCl, 30 mM trisodium citrate, pH 7.0) and the samples were finally rinsed with and stored in the same buffer without DAPI prior to the addition of the photostabilising imaging buffer (PBS containing an oxygen scavenging system of 2.5 mM protocatechuic acid, 10 nM protocatechuate-3,4-dioxygenase supplemented with 1 mM (±)-6-hydroxy-2,5,7,8-tetramethylchromane-2-carboxylic acid (Trolox)^87^.

TagPAINT imaging^88^ was carried out for SNAP-tagged NSP2-expressing cells infected with RVs. After fixation, cells were permeabilized with PBS supplemented with 0.2% Triton-X100 for 3 min, and then subsequently incubated with 50 mg ml^−1^ BSA in PBS for 10 min. 5 μM benzylguanine (BG)-conjugated DNA (Biomers.com) dissolved in PBS supplemented with 0.2% Tween-20 (PBST) was incubated with fixed cell samples for 15 min. The samples were then washed with 0.4 ml of PBST several times, to remove any non-specifically adsorbed ligand. Finally, the samples were incubated with gold nanoparticles in PBST for 10 min before mounting for DNA-PAINT imaging.

### DNA-PAINT Imaging

#### Microscope configuration

DNA-PAINT imaging was carried out on an inverted Nikon Eclipse Ti microscope (Nikon Instruments) equipped with the Perfect Focus System using objective-type total internal reflection fluorescence (TIRF) configuration (oil-immersion Apo SR TIRF, NA 1.49 100x objective). A 200 mW 561 nm laser beam (Coherent Sapphire) was passed through a clean-up filter (ZET561/10, Chroma Technology) and coupled into the microscope objective using a beam splitter (ZT561rdc, Chroma Technology). Fluorescence light was spectrally filtered with an emission filter (ET575lp, Chroma Technology) and imaged with an sCMOS camera (Andor Zyla 4.2) without further magnification, resulting in an effective pixel size of 130 nm after 2×2 binning. Images were acquired using a region of interest of 512×512 pixels. The camera read-out rate was set to 540 MHz, and images were acquired with an integration time of 200 ms.

#### Sample preparation, imaging and data analysis

5’-ATACATTGA-Cy3B-3’ (Metabion) was used as ssDNA ‘imager’ for visualising Seg 3 RNA target. 20,000 frames were acquired for each target. Sequences of the oligonucleotide RNA FISH probes are listed in Supplementary File 1). These were generated using the Stellaris RNA FISH probe designer (https://www.biosearchtech.com/stellaris-designer), using each gene-specific ORF sequence as inputs and level 2 masking. The resulting pools of probes were then further filtered to remove the sequences targeting the RNA transcripts sequences with higher propensity to form stable intra-molecular base-pairing. 5’-TAATGAAGA-Cy3B-3’ (Metabion) was used as ssDNA ‘imager’ for a complementary benzylguanine (BG)-conjugated oligonucleotide DNA (Biomers.com) for reacting with the SNAP-tagged NSP2. Imager strands were diluted to 100 pM (Seg3 RNA), and 300 pM (SNAP-tagged NSP2), respectively. Drift correction was performed with a redundant cross-correlation and gold particles used as fiducial markers. Fluorescence data were subjected to super-resolution reconstruction using Picasso software package^89,90^.

## Authors’ contribution

A.B, F.G., G.P., W.A., G.K., H.E., T.P.J.K., J.A. designed and carried out experiments, and analyzed data. K.L.S., W.A., N.E., R.Q., G.K., T.P.J.K., R.J., S.S. contributed novel analytical tools. A.B. managed the project. All authors contributed ideas, discussed the results and were involved in writing of the manuscript.

## Supplementary Information

**Supplementary Figure 1.**
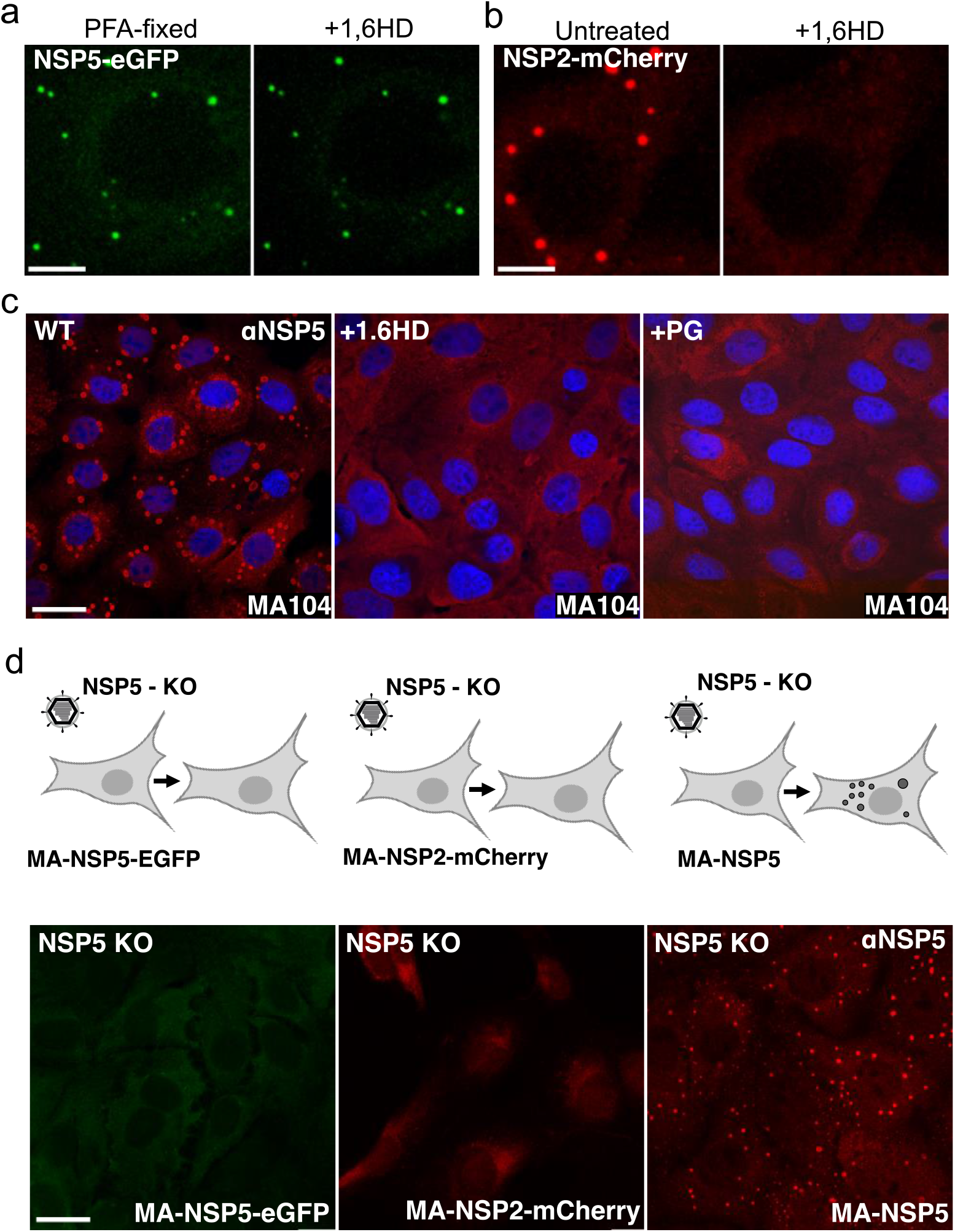
(a) RV-infected MA104-NSP5-EGFP cells (4 HPI) fixed with 4% (v/v) paraformaldehyde for 5 min (PFA-fixed, left panel). Application of 1,6HD (5%) does not dissolve NSP5-EGFP granules after chemical cross-linking with PFA (right). (b) Live-cell images of RV-infected MA104-NSP2-mCherry cells, shown in Fig.1, at 4 HPI. NSP2-mCherry-tagged replication factories dissolve upon application of 4% (v/v) 1,6HD (SI Movie). Scale bars, 10 μm. (c) Immunofluorescent (IF) staining of viral replication factories in RV-infected MA104 cells 6 HPI, before (left), and after a brief (5 min) application of 4% 1,6HD or propylene glycol (PG), respectively, prior to PFA fixation and IF detection of NSP5 (red). Nuclei are stained with DAPI (blue). (d) Recombinant rotavirus NSP5 KO (NSP5 knockout) infection of MA104-derived stable cell lines producing NSP5-EGFP (*left, diffuse* EGFP signal), NSP2-mCherry (*middle, diffuse mCherry signal*), and the wild type NSP5 (*right,* NSP5-rich condensates, IF staining). All cells were fixed and imaged 8 h after infection with NSP5-KO RV. Scale bar, 10 μm.

**Supplementary Figure 2.**
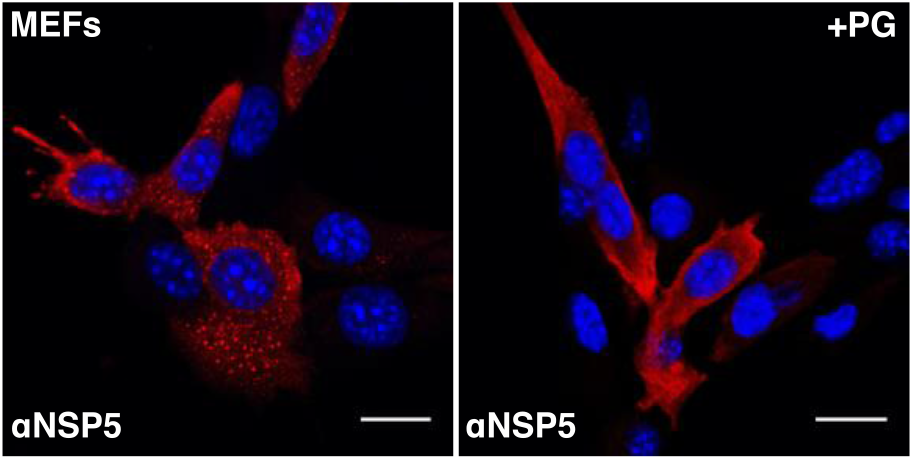
IF staining of NSP5-rich condensates (replication factories) in RV-infected Murine Embryo Fibroblasts (MEFs), 6 HPI. Left – untreated RV-infected MEFs; right – RV-infected MEFs 5 min after treatment with 4% (v/v) propylene glycol (PG). Nuclei – DAPI staining (blue), NSP5 – red. Scale bar 15 μm.

**Supplementary Figure 3.**
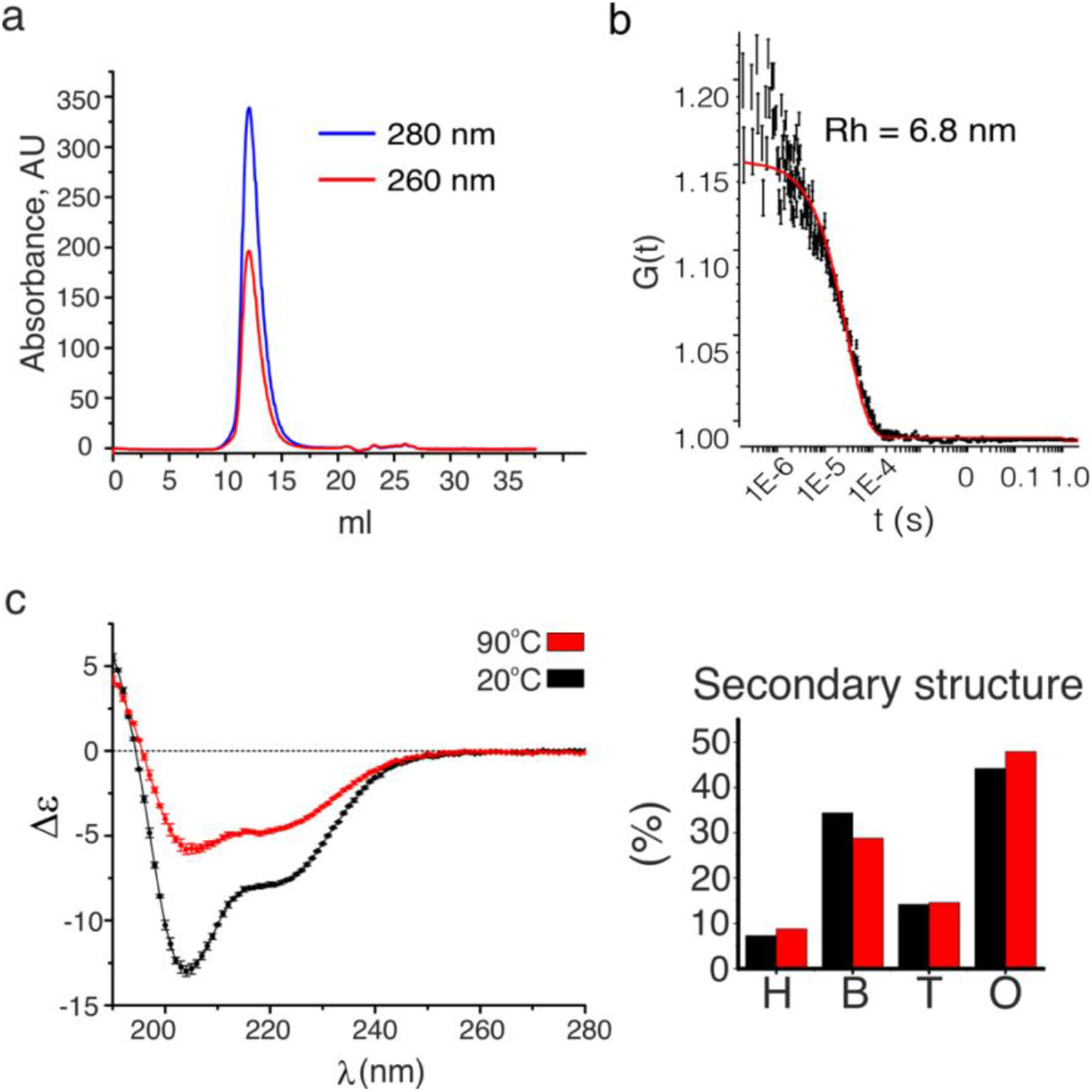
**(a)** Size-exclusion (SEC) analysis (Superdex 200 Increase 10/300 GL) of the purified recombinant protein NSP5. After purification and refolding, the protein was monodisperse and free of nucleic acids, as judged by the A_260_/A_280_ ratio. **(b)** Quasi-elastic scattering analysis of the SEC peak fraction shown in **(a)**, with the calculated hydrodynamic radius, R_h_ ~ 6,8 nm. **(c)** Circular dichroism (CD) spectra of NSP5 acquired at 25°C (black) and after thermal denaturation at 90oC (red). Secondary structure analysis of NSP5 determined by spectral deconvolution of the CD spectra recorded at 25oC (black) and after the thermal denaturation (red). H – helices, B – β-sheets, T – turns, O – disordered.

**Supplementary Figure 4.**
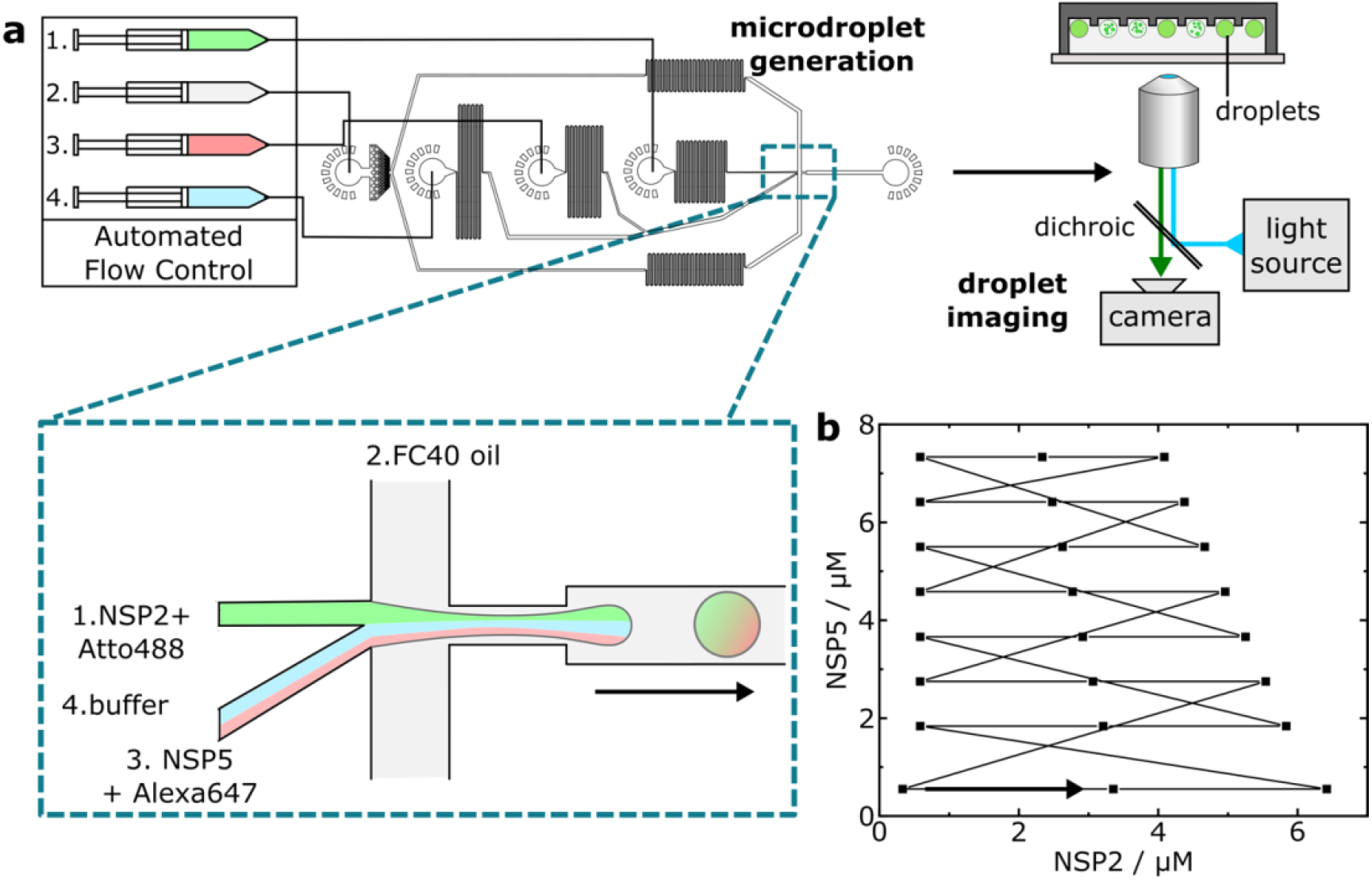
**(a)** Droplets were generated using a microfluidic device controlled by automated syringe pumps. Combination of aqueous droplet components prior to the droplet-generating junction (inset) enables variation in droplet solution composition. Droplets are collected (6 min collection time) off-chip, before undergoing analysis by epifluorescence microscopy. **(b)** Flow profile for NSP2 and NSP5 concentrations as produced by automated flow control in droplet generation. Flow set points (black squares) are maintained for 7 s, with the overall flow programme lasting 168 s. The arrow indicates the beginning of the continuous flow programme loop.

**Supplementary Figure 5.**
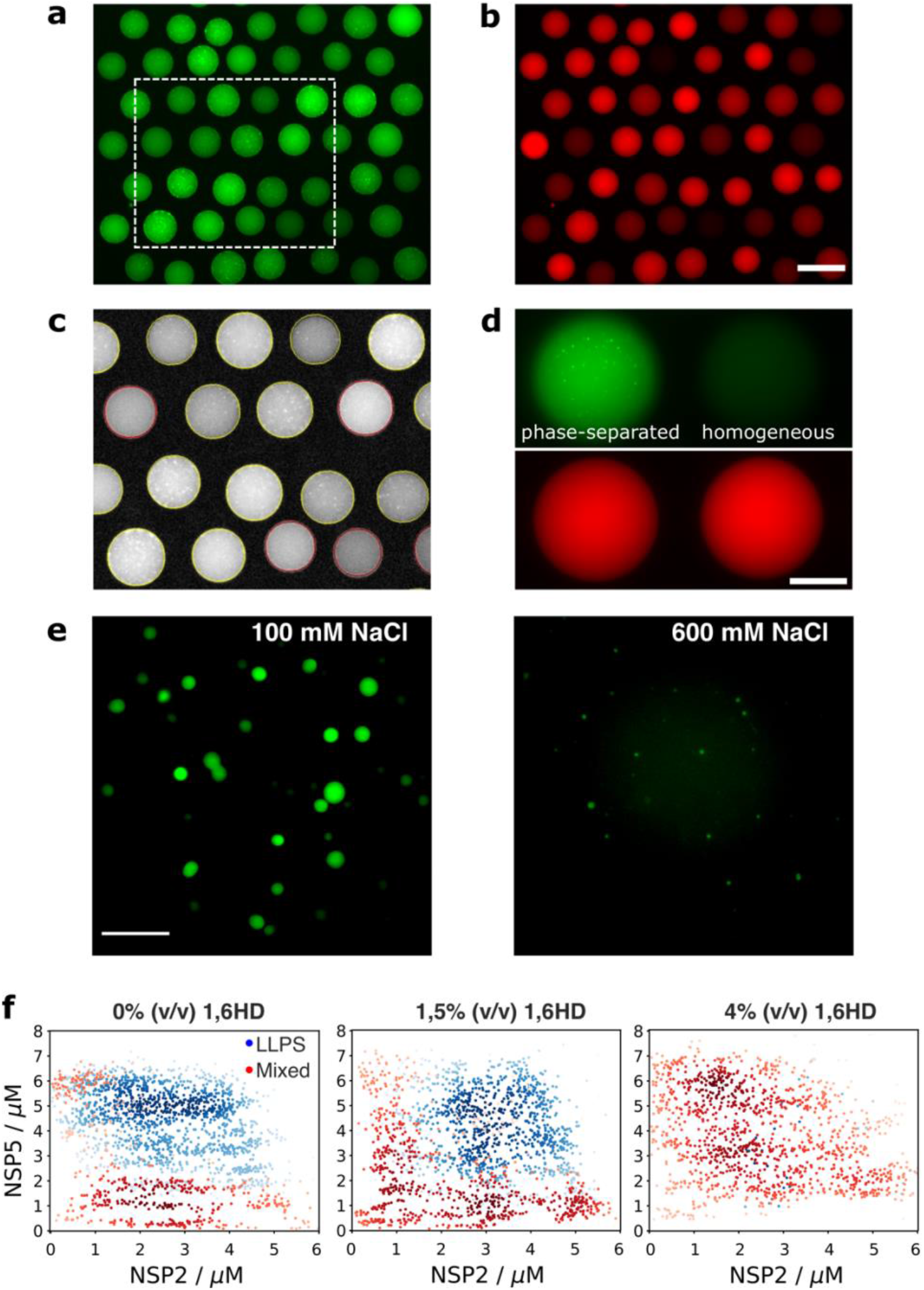
**(a-d)** Representative epifluorescence data for 0% v/v 1,6-hexanediol phase diagram of trapped microdroplets and barcode fluorescence imaged in **(a)** 488 and **(b)** 647 channels. Scale bar = 200 μm. **(c)** Fit of droplet outlines and phase-separation classification output for region enclosed by dashed box in (a), red and yellow outlines denote droplet classification as homogeneous and phase-separated, respectively. **(d)** Representative images of microdroplets and barcode fluorescence classified as phase-separated (left) and homogenous (right) imaged in 488 (upper) and 647 (lower) channels. Scale bar = 100 μm. **(e)** Epifluorescence data for NSP5/NSP2 condensates (5 μM each) formed in the presence of 100 mM and 600 mM NaCl, 488 nm excitation. Scale bar = 10 μm. **(f)** Phase diagrams generated through droplet microfluidics for the coacervation of NSP2 and NSP5, in the presence of 0% v/v (*left*) 1.5% v/v (*middle*) and 4% v/v (*right*) 1,6-hexanediol. Each point corresponds to one droplet microenvironment, with the presence or absence of phase separation denoted by red and blue colouring, respectively. The opacity of each point corresponds to the local density of data for each phase separation classification. Phase diagrams were generated from N = 2206, 2035 and 1470 data points for each 1,6-hexanediol concentrations, respectively.

**Supplementary Figure 6.**
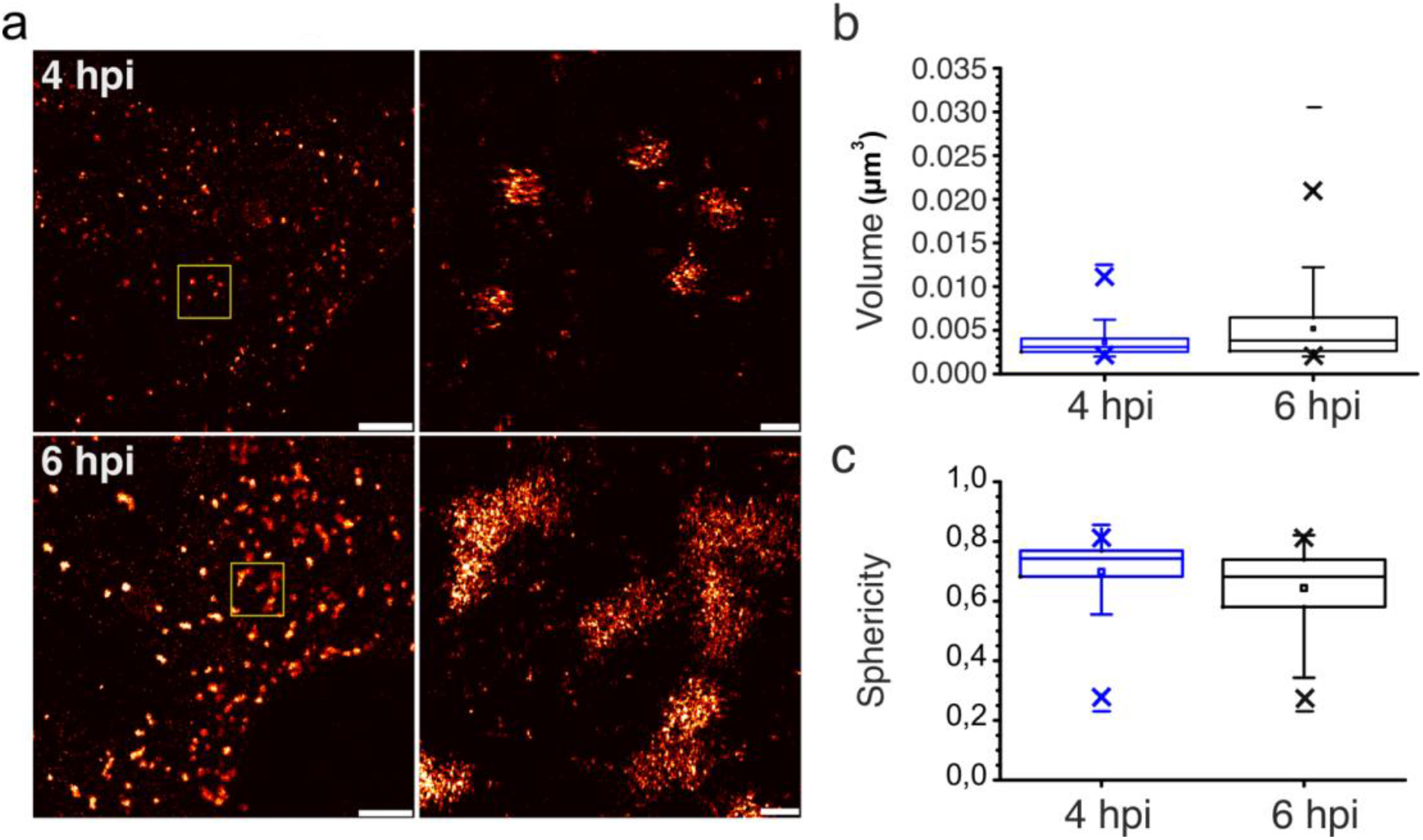
Seg3 RNA foci in RV-infected cells at 4 and 6 HPI imaged through 3D DNA-PAINT. Scale bars, 2 μm (left) and 200 nm (zoomed-in, right). **(b-c)** Distribution of calculated volumes and sphericities of the Seg3 RNA-containing granules in RV-infected cells at 4 HPI (N=704) and 6 hpi (N=698), shown in **(b)**. At the 0,001 level, the two distributions of are significantly different between 4 and 6 hpi, assessed by the two-sample Kolmogorov-Smirnov test.

